# Global and local mechanical properties control endonuclease reactivity of a DNA origami nanostructure

**DOI:** 10.1101/640847

**Authors:** Antonio Suma, Alex Stopar, Allen W. Nicholson, Matteo Castronovo, Vincenzo Carnevale

**Affiliations:** Institute for Computational Molecular Science, Temple University, Philadelphia, PA 19122, USA; Department of Biology, Temple University, Philadelphia PA, 19122, USA; Department of Chemical Science and Technologies, University of Rome, Tor Vergata, Via della Ricerca Scientifica, 00133, Rome, Italy; School of Food Science and Nutrition, University of Leeds, Leeds, UK

## Abstract

We used coarse-grained molecular dynamics simulations to characterize the global and local mechanical properties of a DNA origami triangle nanostructure. The structure presents two metastable conformations separated by a free energy barrier that is lowered upon omission of four specific DNA staples (defect). In contrast, only one stable conformation is present upon removing eight staples. The metastability is explained in terms of the intrinsic conformations of the three trapezoidal substructures. We computationally modeled the local accessibility to endonucleases, to predict the reactivity of twenty sites, and found good agreement with the experimental data. We showed that global fluctuations affect local reactivity: the removal of the DNA staples increased the computed accessibility to a restriction enzyme, at sites as distant as 40nm, due to an increase in global fluctuation. These results raise the intriguing possibility of the rational engineering of allosterically modulated DNA origami.

## INTRODUCTION

DNA origami are two- or three-dimensional nanostructures, resulting from the assembly of hundreds of single-stranded (ss)DNA molecules (staples) with a long ssDNA (scaffold) [1–5]. The ease of design and preparation, and the exquisite control of design at the nanoscale makes DNA origami ideal molecular platforms for applications in biology, medicine, biophysics, and materials science [6–11].

DNA origami provide a unique mechanical material: patterns of soft [single-stranded (ss)] and stiff [double-stranded (ds)] domains can be generated with single nucleotide precision within nanostructures that are able to adopt inter-convertable conformations [12–15]. Moreover, a precise external control of such nucleic acid nanostructures can be attained through a variety of experimental manipulations. Studies have demonstrated control of nanostructures behavior through buffer conditions [16–18], enzyme action with particular attention to endonuclease reactivity [19–23], and electric fields [24–27].

Very recently, our group discovered unexpected allosteric behavior in a DNA triangle [19], a DNA origami first introduced by Rothemund, and subsequently used in several studies [23, 28–30]. Specifically, we investigated the action of several sequence-specific, DNA-cutting enzymes (restriction endonucleases or REases) towards their recognition sites present in the M13 scaffold sequence. We showed that REases act in a binary fashion, as certain sites cannot be cut even at extended reaction times, or, for HhaI REase recognition sites in particular, become reactive following the introduction of a distant structural defect in the triangle (i.e. by omitting, in the self-assembly process, four staples at a distance of approximately 40 nm from the REase site).

In this study, we computationally examine this phenomenon to obtain insight on the global mechanical behavior of the DNA origami triangle and its influence on local parameters such as site reactivity towards restriction endonucleases. We studied three versions of the triangle: the first contains the full complement of staples; the second and the third are triangles deficient in four and eight staples, respectively, in a localized region on the bottom trapezoid. We determined the global mechanical properties of the three triangles, as well as the detailed dynamics of the three trapezoids substructures composing the triangle, and compared their fluctuations with that of the isolated trapezoid. We then correlated these properties with the reactivity of individual sites towards cleavage by HinP1I REase (a neoschizomer of HhaI).

Our findings are as follows. First, the absence of selected staples affects the amplitude of the global conformational fluctuations of the triangle. Two metastable states are observed in the triangle, while omission of four staples lowers the free energy barrier between the two states, and omission of eight staples completely abolishes the barrier, with the free energy profile reducing to a single minimum. The metastability originates from an intrinsic property of the isolated trapezoid, which exhibits two alternate conformations. When coupled into the triangle, the three trapezoids fluctuate synchronously between the two states, thereby giving rise to the metastability of the triangle. Second, we devised a computational method that estimates the steric accessibility of REase sites in DNA origami, and obtained good agreement between the experimental and theoretical results. Specifically, the results reveal that site accessibility depends on local conformational fluctuations. Third, we correlated theoretical site accessibility with the global conformational fluctuations of the triangles and of the isolated trapezoid, thereby showing how endonuclease reactivity in a strongly coupled mechanical system can be modulated in an allosteric fashion.

## MATERIALS AND METHODS

### System setup and simulation details

We performed extensive Langevin molecular dynamics simulations to determine the conformational dynamics of three triangles that differ in staple composition, and of the isolated trapezoid substructure too. The fully intact structure, hereafter referred to as “triangle”, has the scaffold organized by the full complement of staples (Fig. 1a). The second and the third structures lack four staples and eight staples, respectively, in a region that bridges the seam of one of the three trapezoidal elements forming the triangle, and thus they are referred to as “four staple deficient (4sd) triangle” and “eight staple deficient (8sd) triangle” (Fig. 1b and 1c). The fourth structure is an isolated trapezoid identical to those comprising the triangle (Fig. 3a). The first two structures were previously studied experimentally in ref. [19].

**FIG. 1.**
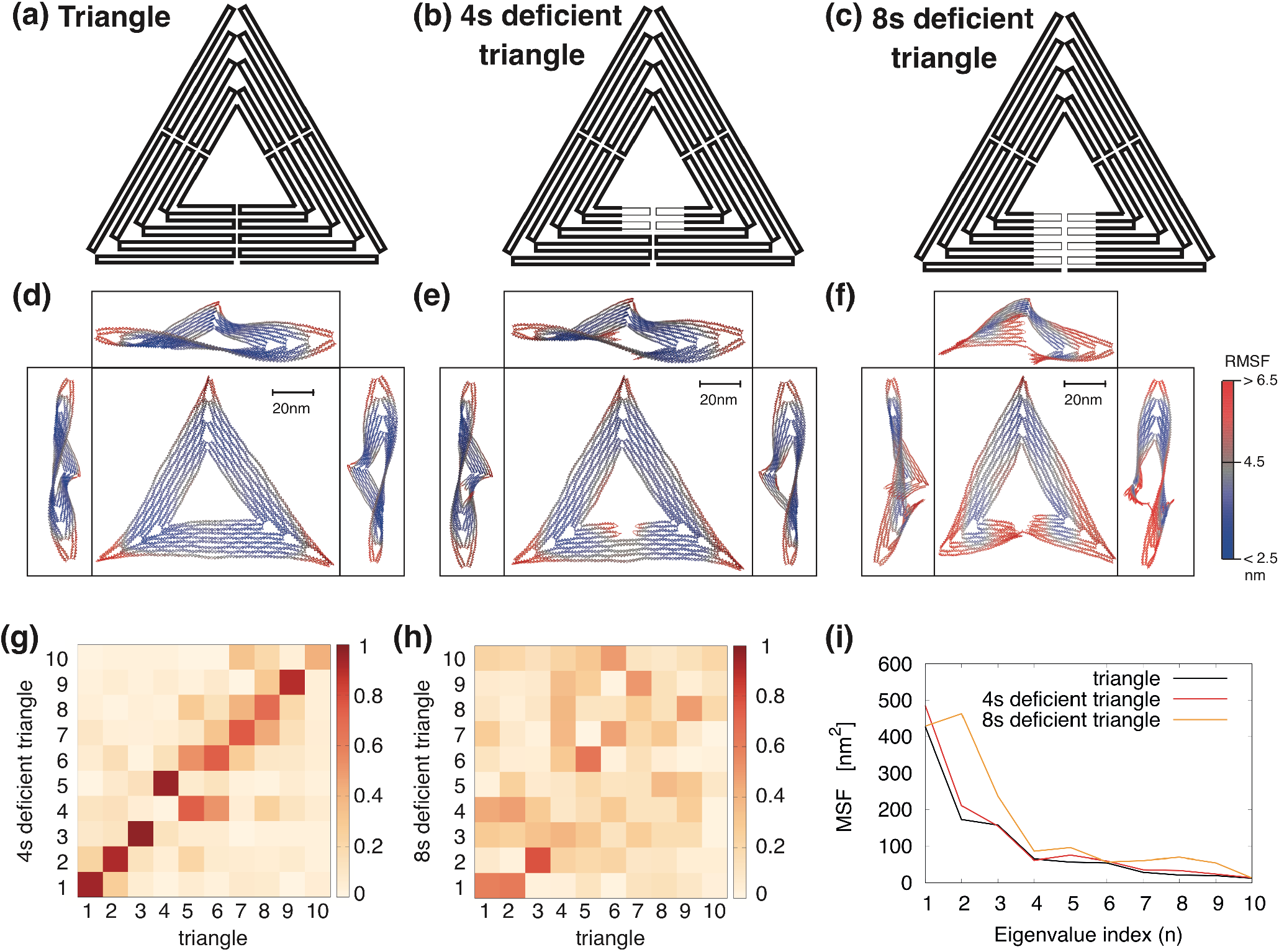
Schematic representation of the scaffold routing map for the triangle (a), the 4sd triangle (b) and the 8sd triangle (c) (sd, staple deficient). The thinner lines in (b)-(c) represent the scaffold portion left unpaired after the removal of four or eight staples, respectively (deficient regions). (d) The triangle, (e) the 4sd triangle and (f) the 8sd triangle average structures computed across the trajectory, viewed from the top and from the three sides. Every trapezoid of the triangle shows a similar S-profile that is visible across the three side views, except for the defective region of 8sd. The color scale indicates the root mean square fluctuation (RMSF) values. Larger fluctuations either concentrate on the tips or are near the defect. (g) Absolute value of the inner product between the first 10 eigenvectors (normalized) of triangle and of 4sd triangle. Except for an inversion between eigenvector 4 and 5, all other eigenvectors have similar directions. (h) Same as (g), between eigenvectors of triangle and of 8sd triangle. Only the first eigenvector is partially conserved. (i) Mean square fluctuations of the three triangles along the directions of the triangle eigenvectors.

The triangles and the trapezoid are composed of about approximately 14000 and 4700 nucleotides, respectively, and were modelled using the second version of oxDNA [31], a coarse-grained DNA model that has been shown to accurately reproduce the mechanical properties of double-stranded and single-stranded DNA molecules [32–38] and that parametrize major and minor grooves and different monovalent salt concentrations.

To obtain the initial configurations, we started from the cadnano [39] constructs of ref [19] (see Supplementary Figures S1-S2-S3-S4), and converted them to the oxDNA representation using the tacoxDNA package [40]. In the case of the triangles, the initial structure presented the three trapezoids in sequential order, thus they were roto-translated to obtain the final geometry, and over-stretched bonds were relaxed using the protocol described in ref. [40].

Simulations were carried out with *T* = 300*K* and at 1*M* monovalent salt concentration. The time of evolution of the four structures was monitored for about 20ms for the triangles, and 10ms for the trapezoid; the mapping between the intrinsic and standard time was previously found to yield *τ_LJ_* ~ 0.7*ns* [41, 42] and the configurations were sampled every 7*μs* (for a total of ~3, 000 configurations). Each simulation was run on 140 processors for a maximum of 1.5 months using LAMMPS [43, 44]. The initial setup files are available as Supplementary files.

### Docking of HinP1I to GCGC sites and accessibility measurement

We developed a quantitative method to measure the accessibility of HinP1I REase towards its recognition sequence (GCGC) in the DNA origami. The first step consisted of superimposing an ideal oxDNA B-DNA helix of 10 bp onto the atomistic DNA structure bound to HinP1I in the PDB structure (PBD code 2FL3, [45]), using a rigid body motion [46]. The ideal double helix was then superimposed onto each GCGC sequence of the triangle, for each frame of the trajectory. The same transformation was applied to the protein.

To determine whether the protein docked to a restriction site overlaps with an adjacent DNA strand, two distance maps were built and compared. These maps, *max_i,θ_* and *min_i,θ_* are functions of the site base-pairs (*i* = 1, … 10) and of the angle around each of them *θ* ∈ [−*π, π*], see Supplementary Figure S5 for more details. The maximum distance map, *max_i,θ_*, reports the distance between the site and the protein particle that is farthest from it. The minimum distance map, *min_i,θ_*, reports the distance between the site and the closest particle from an adjacent DNA strand. Clearly, if *max_i,θ_* < *min_i,θ_* for every *i* and *θ*, the enzyme satisfies the accessibility requirement and the configuration is classified as “accessible”.

## RESULTS

### Global mechanical properties and metastable conformations of the DNA triangle

For a first characterization we computed the average structure and the root mean square fluctuation (RMSF) of each nucleotide around the mean position for the three triangles, see Fig. 1d-e-f. To this end, we applied optimal rotations and translations (so as to minimize the root mean square deviation with respect to the initial configuration) to each frame of the trajectory using the Kabsch algorithm [46].

For all triangles, the average structure is globally twisted, due to the unnatural helical pitch of 10.67 bp/turn imposed by the crossover structures [1, 47]. The fluctuations, highlighted in red, are mostly concentrated at the vertices that exhibit an RMSF greater than 6.5 nm and also near the defects in the 4sd and 8sd triangles.

At first glance, the triangle and the 4sd triangle do not exhibit evident differences that might rationalize the experimental results of ref. [19], except for the area bearing the defect and the edges of the trapezoids, which show slight variations. A deeper analysis of the trajectories, however, show that the triangles fluctuate in different ways over time (see Movies S1-S2-S3).

To distinguish between the different global fluctuation modes, we performed principal component analysis [48] over the trajectories using GROMACS [49]. We constructed the covariance matrix for the three different triangles as 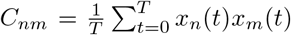, where t is the frame index and *x_n_* is the n-th component of the 3N-dimensional vector of deviations from the average position (*n* = 1, … 3*N* are the coordinates of the scaffold only, in order to compare fluctuations across triangles using the same number of particles), and we computed eigenvalues and eigenvectors ({*λ_n_*, **p**_*n*_}, 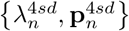 and 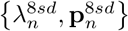) for the triangle, the 4sd and 8sd triangles, respectively.

We projected the first 10 normalized eigenvectors of the triangle on the first 10 ones of the 4sd and 8sd triangles, 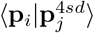 and 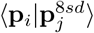 (Fig. 1g-h), and observed a good collinearity for *i* = *j* between triangle and 4sd triangle eigenvectors (except for the inversion between eigenvector 4 and 5), while for the 8sd triangle only the first eigenvector shows good collinearity. Thus, we can quantitatively compare the mean square fluctuations of the triangle and 4sd triangle along the triangle eigenvectors (Fig. 1i), and observe that removal of 4 staples perturbs mainly the first mode, increasing the fluctuation amplitude of 101*nm*^2^. In contrast, removal of 8 staples heavily affects the directions of the principal modes, thus a precise comparison could not be made.

To further analyze the first fluctuation mode, we projected the trajectories over **p**_1_ (see Fig. 2a). This allowed us to analyze the fluctuations of the first mode as a function of time. What emerges is the existence of two metastable conformations, with transitions observed in the triangle and 4sd triangle. In particular, we observe that the triangle resides in each metastable state much longer than the 4sd triangle, while for the 8sd triangle there are no metastable states. By reconstructing the free energy profiles as a function of ⟨**x|p**_1_⟩ (Fig. 2b), we estimate for the triangle an energy barrier between the two minima of about 5*k_B_T* and 2*k_B_T* going from the left minima to the right one and *vice versa*, respectively. For the 4sd triangle, these barriers decrease to about 2.5*k_B_T* and 0.15*k_B_T* . The smaller energy barrier enhances the rate of exchange between the two basins. The 8sd triangle presents only a single minimum.

**FIG. 2.**
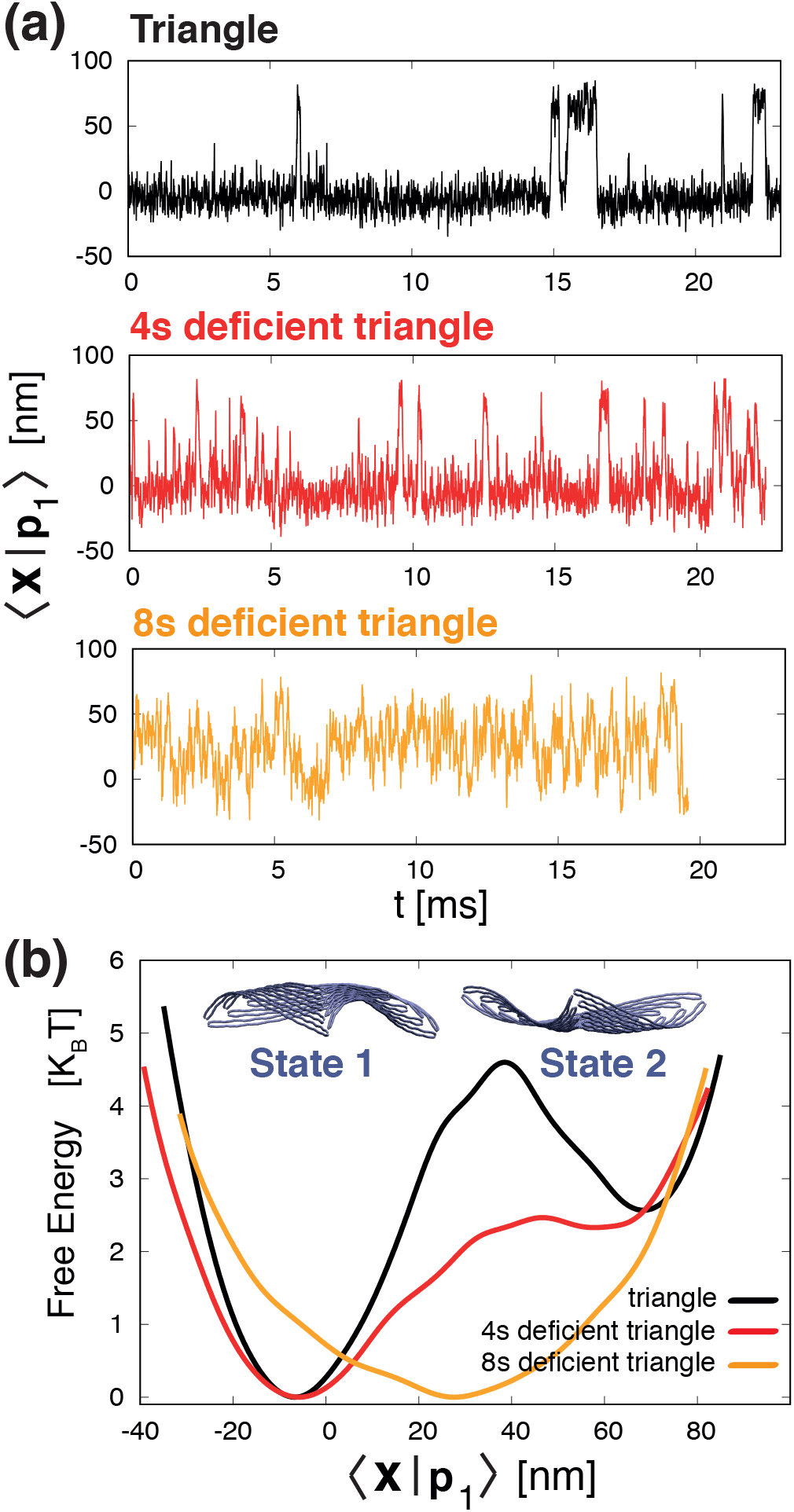
(a) Projections of the trajectories of the triangle and 4sd and 8sd triangles over the first eigenvector of the triangle, ⟨**x|p**_1_⟩. In the first two cases the structure hops between two states, but at different rates, while in the 8sd triangle the fluctuations are over a single minimum. (b) Free energy as a function of ⟨**x|p**_1_⟩. The triangle exhibits two low energy states separated by an energy barrier that is lowered upon introducing a defect of 4 staples; only one minimum is present with an 8 staples defect (where the energy barrier was located in the triangle). The metastable states associated with the two minima are displayed on the top. States 1 and 2 are associated with an upward and downward concavity, respectively.

The two dominant metastable conformations of the triangle that are associated with the two free energy minima are shown in Fig. 2b: the triangle in State 1 is concave upward, while in State 2 it is concave downward. Here, the term upward refers to the direction that is normal to the plane containing the triangle, and that points inward relative to Fig. 4a, while the term downward refers to the opposite direction (see also Movie S1-S2, in which State 1 and 2 are colored in blue and red).

### Mechanical properties of the trapezoid in the DNA triangle

To understand the mechanical origin of the two metastable states, we analyzed separately the time evolution of each of the three trapezoids composing the triangles, designated as left (L), right (R), bottom (B), and compared them with the time evolution of the isolated trapezoid (Fig. 3a-b).

**FIG. 3.**
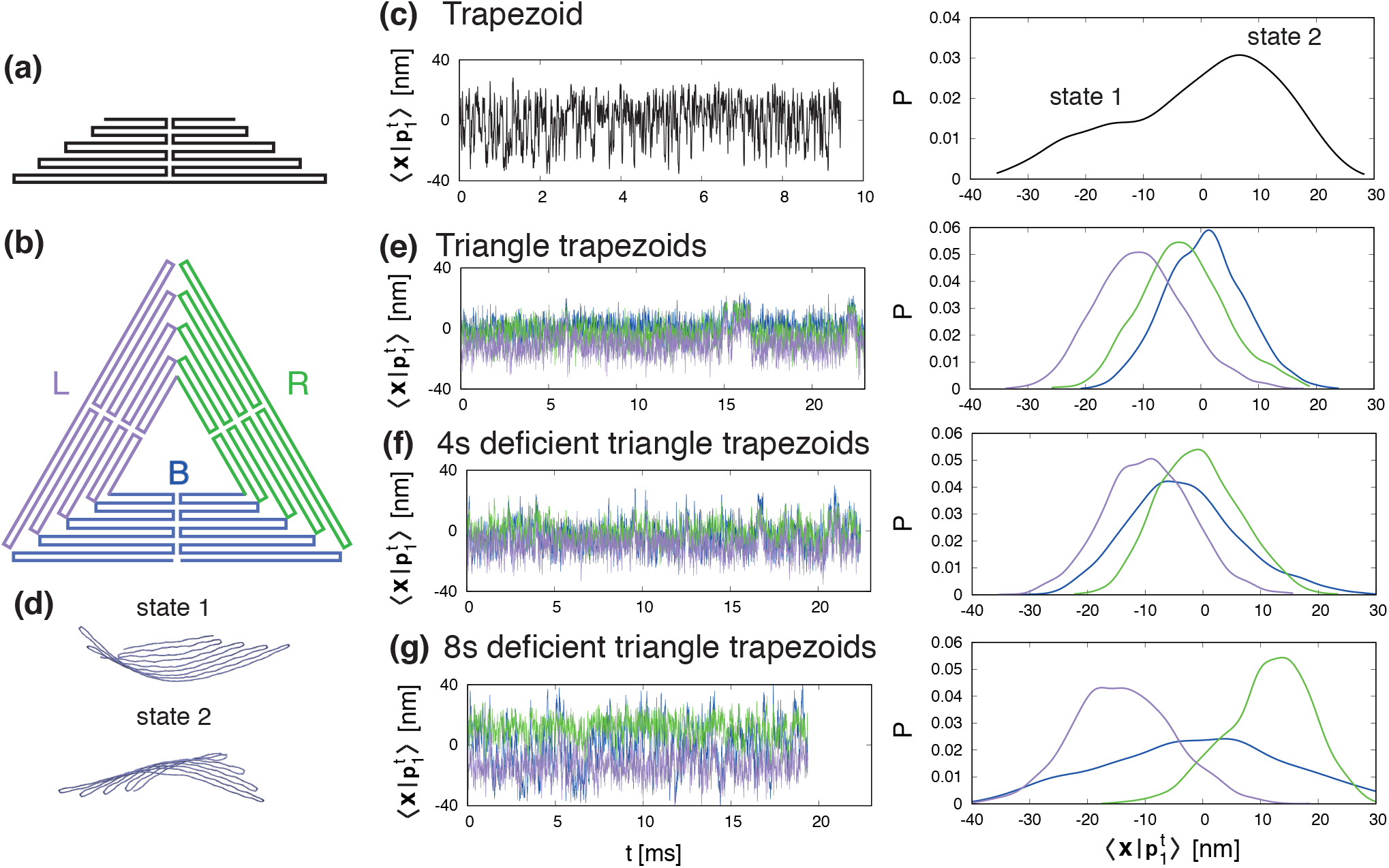
(a) Schematic representation of the scaffold routing map of the isolated trapezoid. (b) Labelling (and associated color) of the three trapezoids composing each triangle: left (L) in purple, right (R) in green and bottom (B) in blue. (c) On the left, projection of trapezoid trajectory along its first eigenvector. On the right, distribution of the projection, showing the existence of two alternate states, depicted in (d). (e-g) On the left, trajectory projection, for each trapezoid substructure composing the three triangles (L,R,B), along the eigenvector of the isolated trapezoid. On the right, distributions of these projections. The line color corresponds to the one defined in (b). In the triangle, the three trapezoids oscillate synchronously between the two trapezoid states. The oscillations correspond to those of the triangle (Fig.2a). The same oscillations appear in the 4sd triangle trapezoids, while in the 8sd triangle the oscillations are decoupled. At the same time, the R-trapezoid distribution peak moves over state 2 while increasing the defect size, and the distribution of the B-trapezoid broadens.

We performed a principal component analysis for each trapezoid (considering only the scaffold), producing a set of eigenvalues and eigenvectors. In particular, the first and most relevant eigenvector of the isolated trapezoid (*p^t^*) is naturally conserved across all the triangle trapezoids (Fig. S6), and thus can be used to project all the trajectories to analyze their fluctuation.

The first eigenvector of the isolated trapezoid describes the transition between two alternate conformations (Fig. 3c-d), differing in their concavity with respect to the surface normal (with state 1 less probable than state 2). When three trapezoids are embedded in the triangle, the geometrical constraints restrict the fluctuations of the three trapezoids between these two states, as evident from the probability distributions of 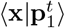 (Fig 3e). Moreover, the fluctuations are highly correlated, and synchronized diffusion between state 1 and state 2 correspond to jumps across the two global metastable states of the triangle. In the 4sd triangle, a similar situation occurs (Fig 3f), but the distribution is broadened by the presence of a defect in the B-trapezoid, a feature that ultimately results in a lowering of the free energy barrier seen in Fig. 2b. In the 8sd triangle the defect is so large that the trapezoids exhibit uncoupled oscillations. Notably the L-trapezoid is preferentially in state 1, while the R-trapezoid is primarily in state 2. This reflects the fact that the L- and R-trapezoids are coupled from two sides, with only the the defective trapezoid side being uncoupled.

### Theoretical accessibility of REase recognition sequences

We developed a method to determine the local accessibility of REase recognition sequences in the triangle. We created a structural proxy by docking the crystallographic structure of the REase to the DNA site under study, then evaluated, for each time frame, the degree of volume overlap between the enzyme and the adjacent DNA structures. A site is scored as accessible if no clash occurs between the two volumes, and inaccessible otherwise (see Fig. 4b and methods for the exact procedure). Finally, we defined the theoretical site accessibility as the fraction of time frames in which the site is accessible to the enzyme.

**FIG. 4.**
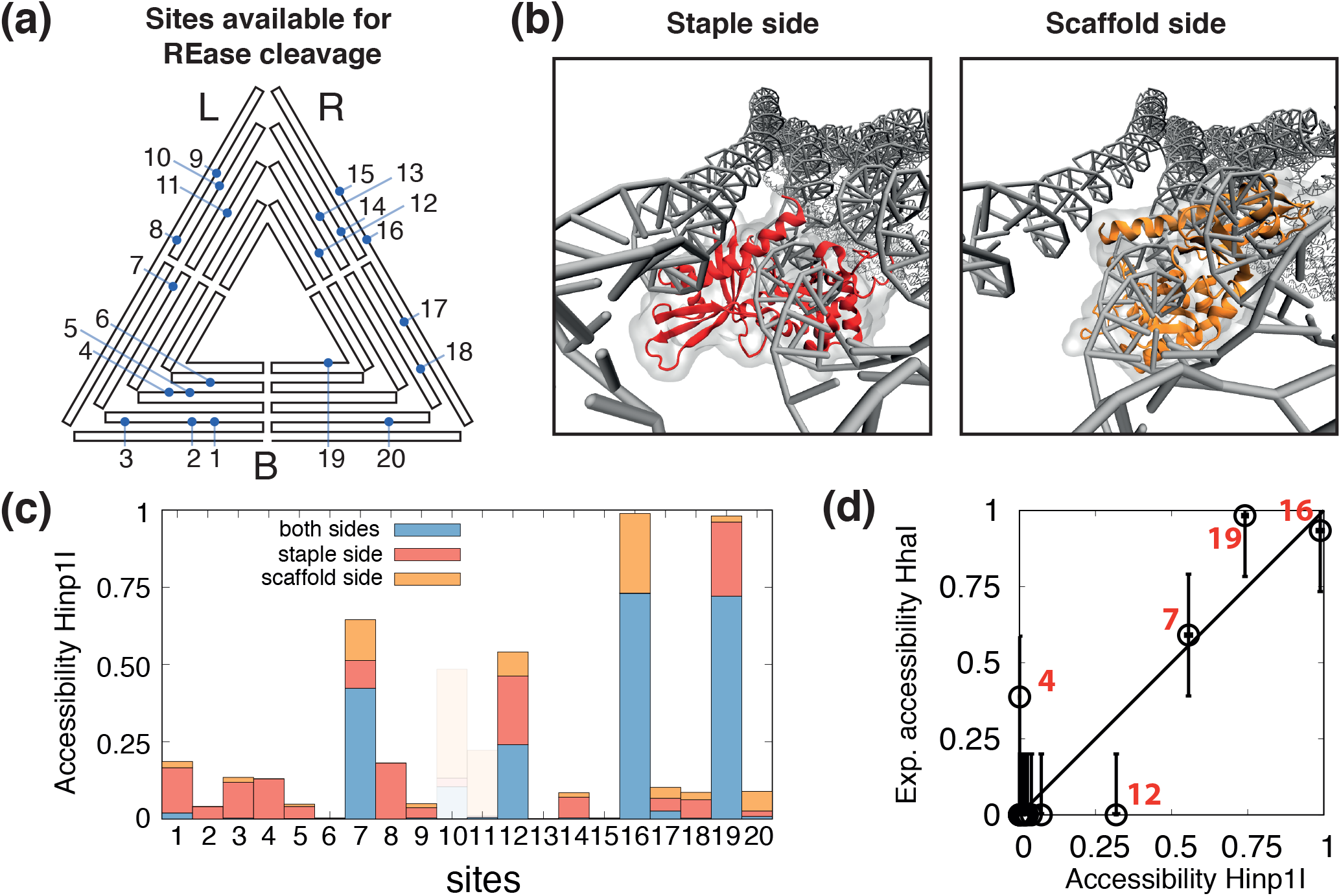
(a) Mapping of the 20 GCGC sequences present in the sharp DNA triangle. b) HinP1I REase docking to site 8 from the staple (red) and scaffold (orange) strand side. (c) Histogram showing the theoretical HinP1I accessibility to the GCGC sequences of the triangle. The accessibility is distinguished by whether the enzyme can access the site only on the scaffold strand (orange), the staple strand (red), or on both strands simultaneously (blue). Sites 10, 11 and 15 (shadowed) were excluded as they overlap with a crossover structure and therefore are not substrates for the enzyme. (d) Scatter plot comparing the theoretical and experimental accessibility from the scaffold strand side. The experimental accessibility is estimated from the gel band intensities as determined in ref. [19] (see Fig. S7). The Pearson’s correlation coefficient is 0.9.

Since the HhaI REase used in ref. [19] lacks a PDB structure, we used HinP1I as a proxy enzyme, as it recognizes the same sequence (GCGC), has a similar mass as HhaI (HinP1I, 28.7 kDa; HhaI, 27.8 kDa), and a crystal structure of HinP1I docked to a duplex DNA has been reported (PDB 2FL3 [45]). Note that HinP1I functions as a monomer, and cuts each strand of the GCGC duplex in separate events [45].

The triangle has twenty GCGC sites (see Fig. 4a), with each exhibiting a scaffold side and a staple side. We distinguished three situations, in which HinP1I can access (i) only the scaffold strand, (ii) the staple strand, or (iii) both strands simultaneously (see Fig. 4c). We excluded sites 10, 11 and 15 (shaded) as they overlap with a DNA crossover junction and therefore are inherently unreactive.

The resultant picture is that four sites exhibit high accessibility : 12, 7, 19, 16 (in ascending order).

We can quantitatively correlate theoretical accessibility to experimental reactivity by proposing that the former quantifies the fraction of reactive triangle conformations. The enzymatic reaction for a specific site *i* can then be described as:

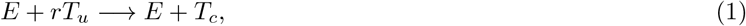

where *E* is the enzyme, *T_u_* are the uncut triangles at site *i*, *r* is the fraction of accessible *T_u_* triangles (0 < *r* < 1), *T_c_* are the cleaved triangles. For a given concentration of *T_u_*, each site *i* will be accessible to the enzyme with a different concentration fraction *rT_u_*. This will be reflected in the kinetics of cleavage: the more accessible sites (with a larger *r*) will be cut more rapidly [50].

We define the experimental accessibility as the fraction of triangles cleaved at a single time point, before the reaction at the most reactive site reaches completion. Figure 4d shows the comparison between the theoretical accessibility for HinP1I and the experimental accessibility, obtained from gel electrophoretic data for a one hour reaction by HhaI [19] (see also Fig. S7). For both sets of data we considered only the accessibility of the scaffold strand, as the experimental data provide information only for that side.

We observed that the theoretical accessibility matches the experimental accessibility within an error margin of 20%, a value comparable to the statistical background “noise” of the gel electrophoretic data. In particular, site 7 is cut more slowly than sites 16 and 19, as predicted by the model. These results support the assumption that the reactivity of a given GCGC site is related to its accessibility. Only sites 4 and 12 are an exception, for multiple possible reasons. Firstly, the use of monovalent instead of divalent cations (implicit in oxDNA) might locally alter the modeled structure, thereby resulting in subtle changes in helix-helix distances [51] and relative orientation. Secondly, the experiments in Fig. S7 were carried out by creating base mismatches involving the two central bases of all the GCGC recognition sites (thus masking the sites) except the site under investigation. These modifications, while blocking REase action, can potentially alter the local fluctuations of neighboring sites, especially near sites 10, 11 and 15 which overlap with crossover structures. Thirdly, sub-nanometric structural differences between HhaI and Hinp1I enzymes may be relevant for specific docking angles. Lastly, there might be a threshold of experimental accessibility, above which each site can be scored as experimentally reactive.

### Local mechanical fluctuations affect endonuclease accessibility

To obtain insight on the local factors that influence site accessibility, we projected the center of mass of the adjacent DNA double-helical structures over a plane perpendicular to the GCGC helical axis, using as a reference frame the direction of one of the GCGC nucleotides (see fig. 5a). Figure 5b shows the projection for three representative sites, that also distinguishes between scaffold and staple sides: site 6, non-accessible from both sides; site 7, accessible from both sides; and site 8, accessible only from the staple side.

**FIG. 5.**
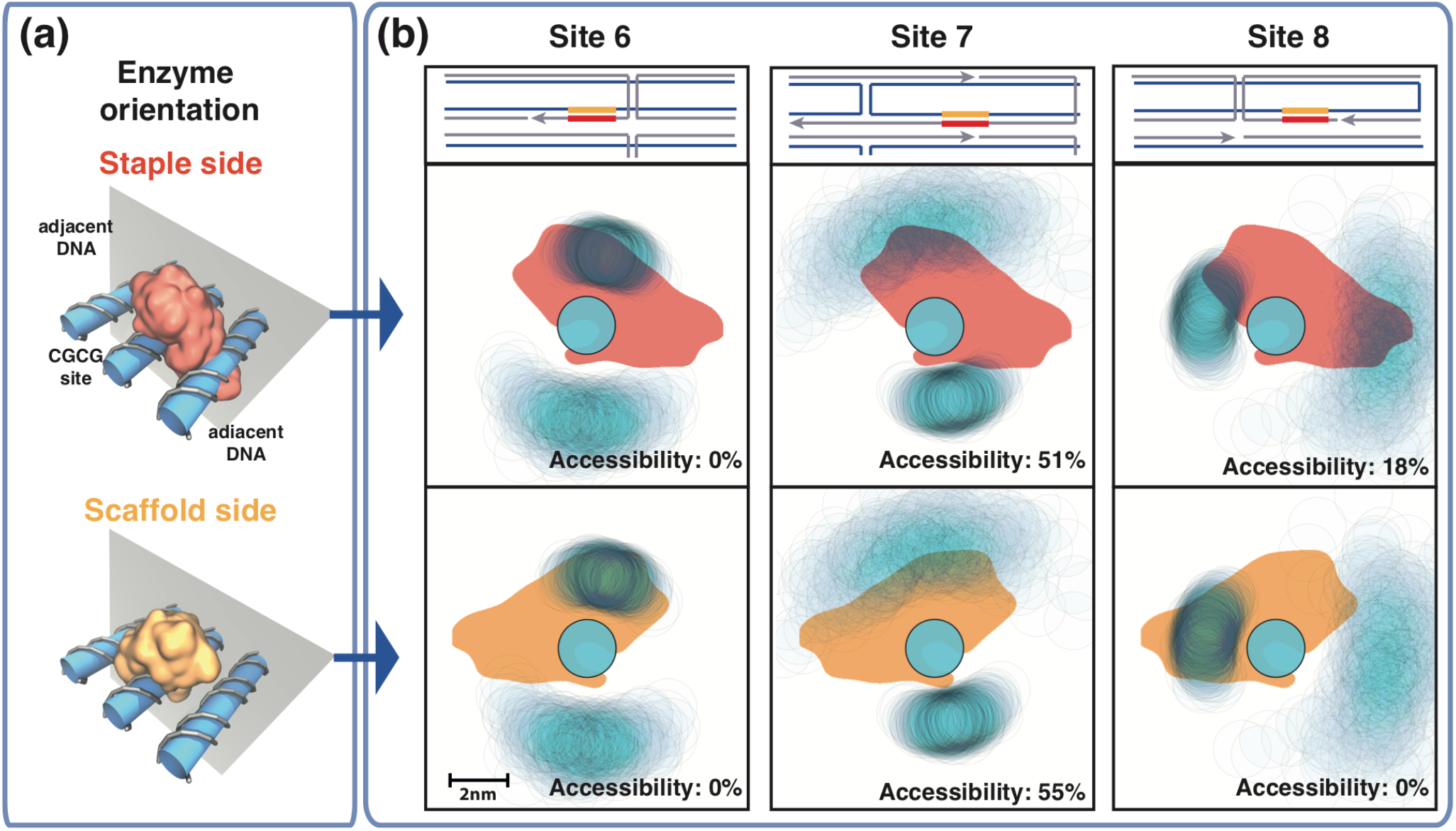
(a) Representation of HinP1I enzyme docking onto the GCGC site in DNA origami, either from the staple strand side or from the scaffold strand side (top and bottom panel, respectively). The adjacent DNA helices are also shown. (b) Top row, local arrangement of the DNA structures surrounding sites 6, 7, 8. The scaffold (staple) strand is colored blue (gray); two close parallel lines denote a DNA double helix, and orthogonal lines describe the path of crossover junctions. Middle (bottom) row, projections of the enzyme docked on the GCGC site from the staple (scaffold) side, colored in red (yellow); of the center of mass of the DNA helix bearing the GCGC site and of the two adjacent DNA helices (in light blue, with a diameter of ~2nm), for sites 6, 7 and 8. The reference frame is the position of one nucleotide of the GCGC site; thus, the positions of the enzyme and the GCGC site are fixed, while the position of the adjacent DNA helices fluctuate, forming clouds with intensities proportional to the probability that a given area is occupied by the DNA helix. For each projection, the theoretical accessibility is reported in the panel. For site 6, the enzyme always overlaps with the upper adjacent DNA helix, resulting in inaccessibility from both sides. For site 7, the enzyme overlaps half of the time with the upper DNA helix from both sides; thus, it is partially accessible in both cases. For site 8, the enzyme overlaps with the left adjacent DNA structure, thus creating inaccessibility only from the scaffold side.

The local accessibility is determined by the degree of overlap between the HinP1I enzyme projection and the adjacent DNA molecules. Complete overlap prevents cleavage, while partial overlap affords accessibility, with a cleavage rate proportional to the percentage of overlap over time. In particular, for site 6, we find that the enzyme always overlaps with the upper adjacent DNA molecule, thus suppressing the site accessibility. For site 7, the enzyme overlaps only 50% of the time with the upper adjacent DNA molecule, due to the higher flexibility of that molecule. For site 8, we find the enzyme overlap with the left adjacent DNA molecule is the most important to suppress site accessibility from the scaffold side, but the same does not apply from the staple side.

We note that the degree of overlap between the enzyme and the adjacent DNA molecules depends on several variables, such as the orientation of the bases in the GCGC sites (from which the orientation of the docking protein depends), and how much the adjacent DNA molecules fluctuate, which in turn is dictated by the distance from a crossover junction.

### Global mechanical fluctuations affect endonuclease accessibility in an allosteric manner

We next compared the local site reactivity between the three triangles and the isolated trapezoid, in order to explore its possible connection with changes in global mechanical fluctuations (Fig. 6). To this aim, we mapped the restriction sites of the triangle onto the equivalent positions on the trapezoid. Qualitatively, we observed that the most remarkable accessibility variation across the studied structures is seen for sites 4 and 8 from the staple side (13% and 18% increase, respectively), and for site 7 from the scaffold side (20% increase).

**FIG. 6.**
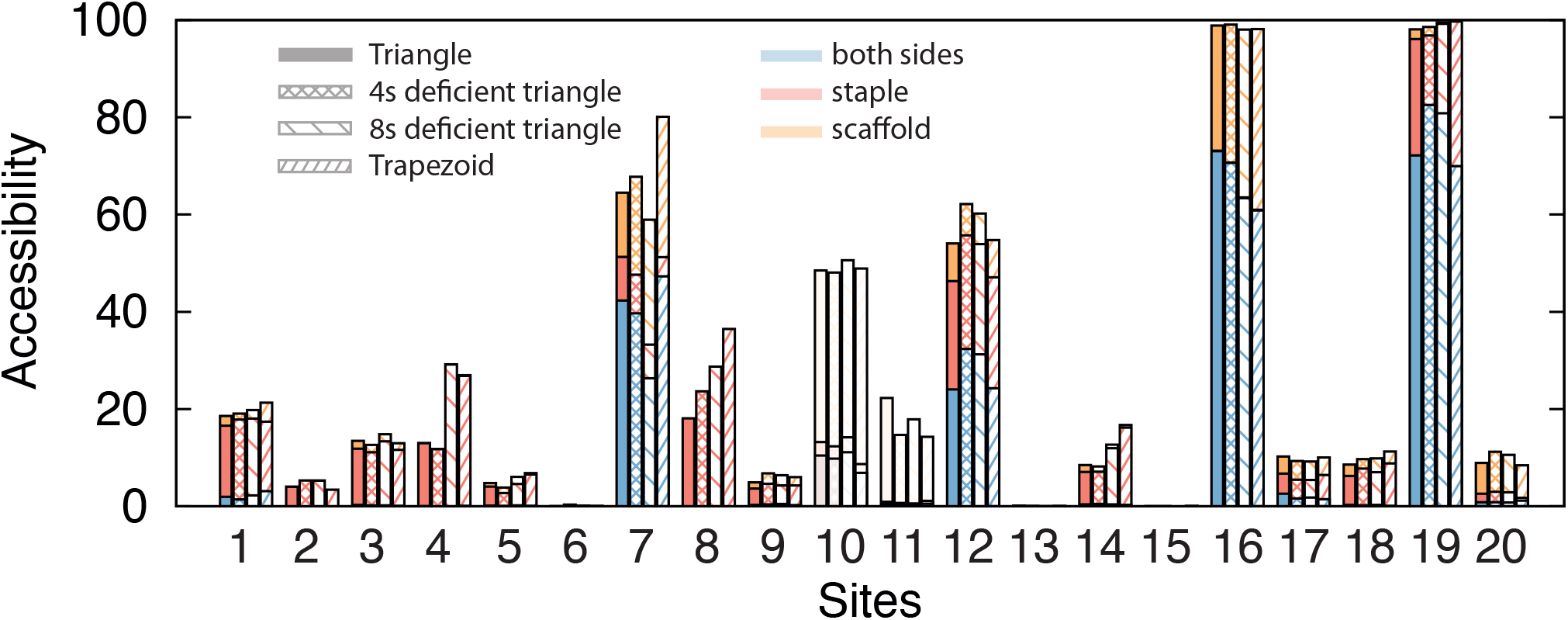
Side-by-side comparison of the theoretical accessibilities of the 20 GCGC sequences for the triangle (solid), 4sd triangle (square mesh), 8sd triangle (coarsely-spaced line mesh), isolated trapezoid (finely-spaced line mesh) (see Fig. 4c for more details). The most consistent increase in accessibility across all sites is for sites 4, 8 from the staple side and site 7 from the scaffold side.

Sites 7 and 8 happen to be in a region of the triangle about 40nm far from the defective region, and far from the border connected to other trapezoids in the case of the single trapezoid. We therefore examined whether there is a correlate of the increase in accessibility, which is mainly associated with local fluctuations (fast modes), to global fluctuations (slow modes). We observed that fluctuations across different eigenvectors are highly correlated to each other (e.g. Fig. S8). For this reason, we speculate that an increase in fluctuation over the slower modes can propagate over the faster ones. This is corroborated by the fact that the total mean square fluctuation (i.e. the sum of all the eigenvalues) of the trapezoids containing site 7 and 8 shows the same trend as its accessibility (Fig. 7).

**FIG. 7.**
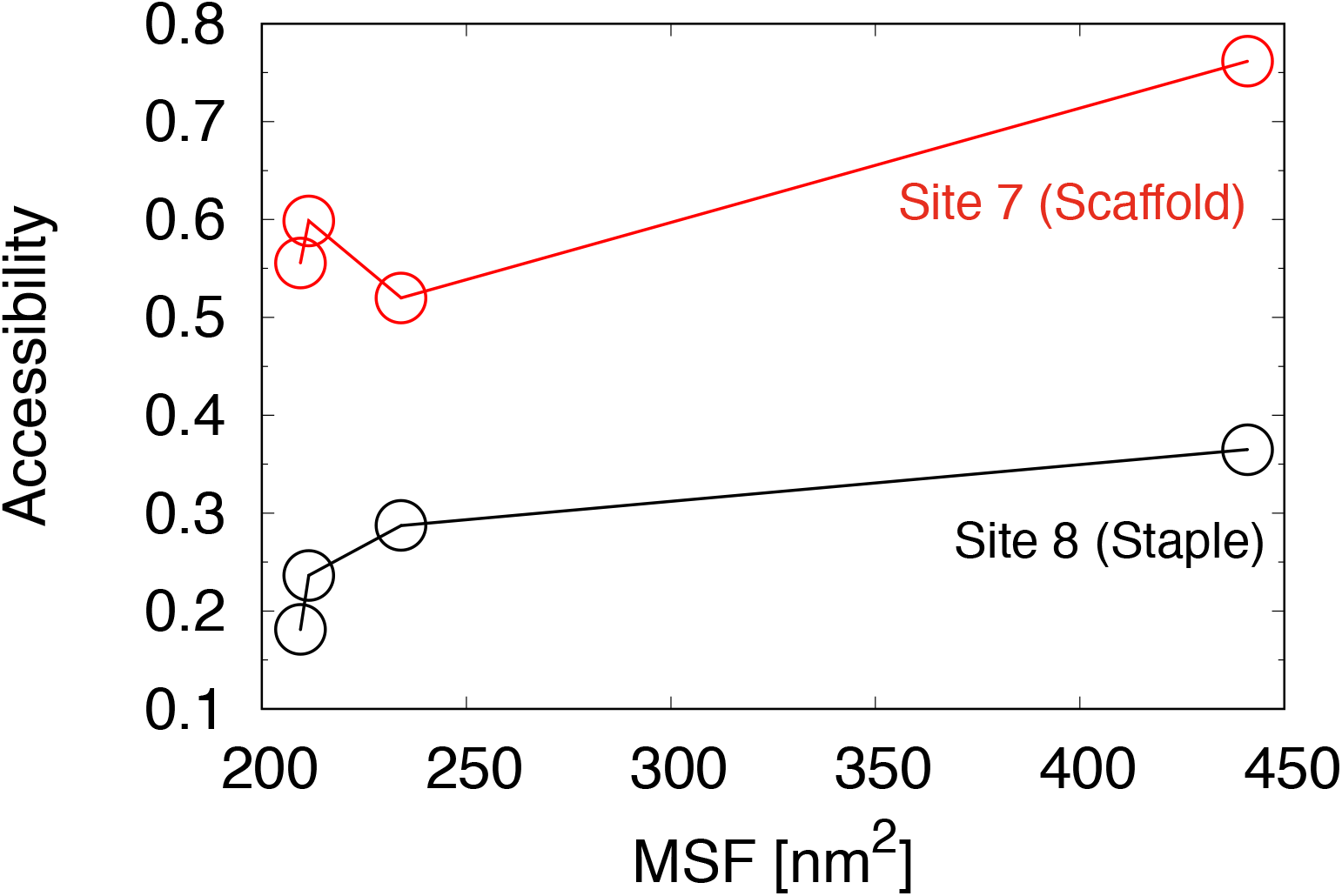
Plot of the side-accessibility of site 7, from the scaffold side, and of site 8 from the staple side, versus total mean square fluctuations (summing over all eigenvalues) for the L-trapezoid of the three triangles (which contains both sites 7 and 8) and the isolated trapezoid.

These results suggest that allosteric effects may be operative in the DNA triangle: while the GCGC sites 7 and 8 are distant from the B-trapezoid, a structural change (staple omission or addition) in the latter can modulate the reactivities of site 7 and 8 by altering its dynamics. In the experiments of ref [19] it was observed that sites 9, 12 and 13 become reactive in the 4sd triangle. Although only site 12 exhibits an increase in local accessibility in the simulations, multiple other factors mentioned above, and not accounted in the model, can contribute to a change in accessibility along with the observed change in global fluctuations.

## DISCUSSION

DNA nanostructures, including DNA origami, constitute a novel class of materials with emergent properties and behaviors yet to be fully understood and applied. Here, we used extensive coarse-grained molecular dynamics simulations to probe the global and local mechanical properties of a triangular DNA origami.

We found that two metastable conformations are present, with the triangle capable of assuming either an upward or downward concave shape with respect to the triangle surface normal. Upon omission of four staples, the interconversion rate between the two states increases, due to a decrease in the intervening free energy barrier. By removing 8 staples over the same defective region, the metastability is lost. These mechanical fluctuations can be understood in terms of mechanical fluctuations of the trapezoid substructures, which intrinsically possess two distinct alternate states. In the triangle, the three trapezoids exchange between the two states in a synchronous manner, resulting in global transitions between the two metastable conformations of the triangle. With the four staple defect, the fluctuations are synchronous, but more frequent. With the eight staple defect, the synchronicity is lost.

We devised a method to quantify the local accessibility of GCGC sequences to a cognate restriction endonuclease, and showed that site reactivity on the scaffold side correlates with the predicted accessibility. In particular, the reactivity of a site is dependent upon the degree of steric overlap between the docked enzyme and the adjacent double helices, which in turn depends on the local structure and conformational fluctuations of the site.

For the three triangles and the trapezoid structures that were analyzed we found that several sites exhibit a positive trend in increased accessibility. These changes in local fluctuations can be in part correlated to changes in global fluctuations, which in general are due to modifications distant from the sites themselves.

This study provides an initial step towards a deeper understanding of global and local mechanical properties of DNA origami, and how these properties can play a role in the control of the DNA accessibility (and reactivity too) to a protein. Future studies intend to explore the relationship between global fluctuation modes and the design arrangement of staples e.g. using a honeycomb lattice, and the quantitative assessment of the kinetic behavior of enzymes acting on these nanostructures (see e.g. see ref [23]). These advancements in turn should enable the rational design of allosteric metamaterials, whose shape and conformational fluctuations are controllable by appropriate localized stimuli [52–55]. Such behaviors can be exploited to develop, for example, carrier platforms with programmable reactivities toward nucleases or with dynamic stimuli-responsive properties.

## ACKNOWLEDGEMENTS

This project received funding from the European Union Horizon 2020 research and innovation programme under the Marie Skodowska-Curie grant agreement No. 645684, a National Institutes of Health Grant S10OD020095 (to V.C.) and the National Science Foundation Grant ACI-1614804 (to V.C.). This research involved calculations carried out using Temple University’s High Performance Computing resources, supported in part by the National Science Foundation through major research instrumentation grant number 1625061, and by the US Army Research Laboratory under contract number W911NF-16-2-0189. We thank Sir Richard J. Roberts for helpful discussions.

## References

[1] Rothemund, P. W. Nature 2006, 440, 297.

[2] Castro, C. E.; Kilchherr, F.; Kim, D.-N.; Shiao, E. L.; Wauer, T.; Wortmann, P.; Bathe, M.; Dietz, H. Nat. Methods 2011, 8, 221.

[3] Yoo, J.; Aksimentiev, A. Proc. Natl. Acad. Sci. USA 2013, 110, 20099–20104.

[4] Douglas, S. M.; Dietz, H.; Liedl, T.; Högberg, B.; Graf, F.; Shih, W. M. Nature 2009, 459, 414.

[5] Seeman, N. C.; Lukeman, P. S. Rep. Prog. Phys. 2004, 68, 237.

[6] Liu, S.; Jiang, Q.; Wang, Y.; Ding, B. Adv. Healthc. Mater. 2019, 1801658.

[7] Ramakrishnan, S.; Ijäs, H.; Linko, V.; Keller, A. Comput. Struct. Biotechnol. J. 2018, 16, 342–349.

[8] Bila, H.; Kurisinkal, E. E.; Bastings, M. M. Biomater. Sci. 2019, 7, 532–541.

[9] Hu, Y.; Niemeyer, C. M. Adv. Mater. 2019, 31, 1970190.

[10] Balakrishnan, D.; Wilkens, G. D.; Heddle, J. G. Nanomedicine 2019, 14, 911–925.

[11] others,, et al. Nature 2018, 559, 593–598.

[12] Dietz, H.; Douglas, S. M.; Shih, W. M. Science 2009, 325, 725–730.

[13] Sharma, R.; Schreck, J. S.; Romano, F.; Louis, A. A.; Doye, J. P. K. ACS Nano 2017, 11, 12426–12435.

[14] Marras, A. E.; Zhou, L.; Su, H.-J.; Castro, C. E. Proc. Natl. Acad. Sci. USA 2015, 112, 713–718.

[15] Andersen, E. S.; Dong, M.; Nielsen, M. M.; Jahn, K.; Lind-Thomsen, A.; Mamdouh, W.; Gothelf, K. V.; Besenbacher, F.; Kjems, J. ACS nano 2008, 2, 1213–1218.

[16] Perrault, S. D.; Shih, W. M. ACS nano 2014, 8, 5132–5140.

[17] Gerling, T.; Wagenbauer, K. F.; Neuner, A. M.; Dietz, H. Science 2015, 347, 1446–1452.

[18] Kielar, C.; Xin, Y.; Shen, B.; Kostiainen, M. A.; Grundmeier, G.; Linko, V.; Keller, A. Angew. Chem. 2018, 130, 9614–9618.

[19] Stopar, A.; Coral, L.; Di Giacomo, S.; Adedeji, A. F.; Castronovo, M. Nucleic Acids Res. 2017, 46, 995–1006.

[20] Agarwal, N. P.; Matthies, M.; Gür, F. N.; Osada, K.; Schmidt, T. L. Angew. Chem. 2017, 56, 5460–5464.

[21] Ponnuswamy, N.; Bastings, M. M.; Nathwani, B.; Ryu, J. H.; Chou, L. Y.; Vinther, M.; Li, W. A.; Anastassacos, F. M.; Mooney, D. J.; Shih, W. M. Nat. Commun. 2017, 8, 15654.

[22] Auvinen, H.; Zhang, H.; Kopilow, A.; Niemelä, E. H.; Nummelin, S.; Correia, A.; Santos, H. A.; Linko, V.; Kostiainen, M. A. Adv. Healthc. Mater. 2017, 6, 1700692.

[23] Ramakrishnan, S.; Shen, B.; Kostiainen, M. A.; Grundmeier, G.; Keller, A.; Linko, V. ChemBioChem 2019, 20, 2818–2823.

[24] Kopperger, E.; List, J.; Madhira, S.; Rothfischer, F.; Lamb, D. C.; Simmel, F. C. Science 2018, 359, 296–301.

[25] Suma, A.; Di Stefano, M.; Micheletti, C. Macromolecules 2018, 51, 4462–4470.

[26] Li, C.-Y.; Hemmig, E. A.; Kong, J.; Yoo, J.; Hernández-Ainsa, S.; Keyser, U. F.; Aksimentiev, A. ACS nano 2015, 9, 1420–1433.

[27] Klotz, A. R.; Soh, B. W.; Doyle, P. S. Phys. Rev. Lett. 2018, 120, 188003.

[28] Ding, B.; Deng, Z.; Yan, H.; Cabrini, S.; Zuckermann, R. N.; Bokor, J. J. Am. Chem. Soc. 2010, 132, 3248–3249.

[29] Zhang, Q.; Jiang, Q.; Li, N.; Dai, L.; Liu, Q.; Song, L.; Wang, J.; Li, Y.; Tian, J.; Ding, B.; Du, Y. ACS nano 2014, 8, 6633–6643.

[30] Gopinath, A.; Miyazono, E.; Faraon, A.; Rothemund, P. W. Nature 2016, 535, 401.

[31] Snodin, B. E. K.; Randisi, F.; Mosayebi, M.; ulc, P.; Schreck, J. S.; Romano, F.; Ouldridge, T. E.; Tsukanov, R.; Nir, E.; Louis, A. A.; Doye, J. P. K. J. Chem. Phys. 2015, 142, 234901.

[32] Ouldridge, T. E.; Louis, A. A.; Doye, J. P. Phys. Rev. Lett. 2010, 104, 178101.

[33] Ouldridge, T. E.; Louis, A. A.; Doye, J. P. K. J. Chem. Phys 2011, 134, 085101.

[34] Snodin, B. E. K.; Schreck, J. S.; Romano, F.; Louis, A. A.; Doye, J. P. K. Nucleic Acids Res. 2019, 47, 1585–1597.

[35] Šulc, P.; Romano, F.; Ouldridge, T. E.; Rovigatti, L.; Doye, J. P. K.; Louis, A. A. J. Chem. Phys 2012, 137, 135101.

[36] Engel, M. C.; Smith, D. M.; Jobst, M. A.; Sajfutdinow, M.; Liedl, T.; Romano, F.; Rovigatti, L.; Louis, A. A.; Doye, J. P. K. ACS Nano 2018, 12, 6734–6747.

[37] Snodin, B. E. K.; Romano, F.; Rovigatti, L.; Ouldridge, T. E.; Louis, A. A.; Doye, J. P. K. ACS Nano 2016, 10, 1724–1737.

[38] Shi, Z.; Castro, C. E.; Arya, G. ACS Nano 2017, 11, 4617–4630.

[39] Douglas, S. M.; Marblestone, A. H.; Teerapittayanon, S.; Vazquez, A.; Church, G. M.; Shih, W. M. Nucleic Acids Res. 2009, 37, 5001–5006.

[40] Suma, A.; Poppleton, E.; Matthies, M.; Šulc, P.; Romano, F.; Louis, A. A.; Doye, J. P. K.; Micheletti, C.; Rovigatti, L. J. Comput. Chem. 2019, 40, 2586–2595.

[41] Suma, A.; Micheletti, C. Proc. Natl. Acad. Sci. USA 2017, 114, E2991–E2997.

[42] Coronel, L.; Suma, A.; Micheletti, C. Nucleic Acids Res. 2018, 46, 7533–7541.

[43] Plimpton, S. J. Comput. Phys. 1995, 117, 1–19.

[44] Henrich, O.; Fosado, Y. A. G.; Curk, T.; Ouldridge, T. E. Eur. Phys. J. E 2018, 41, 57.

[45] Horton, J. R.; Zhang, X.; Maunus, R.; Yang, Z.; Wilson, G. G.; Roberts, R. J.; Cheng, X. Nucleic Acids Res. 2006, 34, 939–948.

[46] Kabsch, W. Acta Crystallogr. A 1976, 32, 922–923.

[47] Ke, Y.; Douglas, S. M.; Liu, M.; Sharma, J.; Cheng, A.; Leung, A.; Liu, Y.; Shih, W. M.; Yan, H. J. Am. Chem. Soc. 2009, 131, 15903–15908.

[48] García, A. E. Phys. Rev. Lett. 1992, 68, 2696–2699.

[49] Hess, B.; van der Spoel, D.; Lindahl, E.; Smith, J. C.; Shirts, M. R.; Bjelkmar, P.; Larsson, P.; Kasson, P. M.; Schulz, R.; Apostolov, R.; Pronk, S.; Pll, S. Bioinformatics 2013, 29, 845–854.

[50] Golinik, M. J. Enzyme Inhib. Med. Chem. 2013, 28, 879–893.

[51] Luan, B.; Aksimentiev, A. J. Am. Chem. Soc. 2008, 130, 15754–15755.

[52] Yan, L.; Ravasio, R.; Brito, C.; Wyart, M. Proc. Natl. Acad. Sci. USA 2017, 114, 2526–2531.

[53] Laskowski, R. A.; Gerick, F.; Thornton, J. M. FEBS lett. 2009, 583, 1692–1698.

[54] Changeux, J.-P.; Edelstein, S. J. Science 2005, 308, 1424–1428.

[55] Bertoldi, K.; Vitelli, V.; Christensen, J.; van Hecke, M. Nat. Rev. Mater. 2017, 2, 17066.

