## Supplementary material for "Global and local mechanical properties control endonuclease reactivity of a DNA origami nanostructure": SI

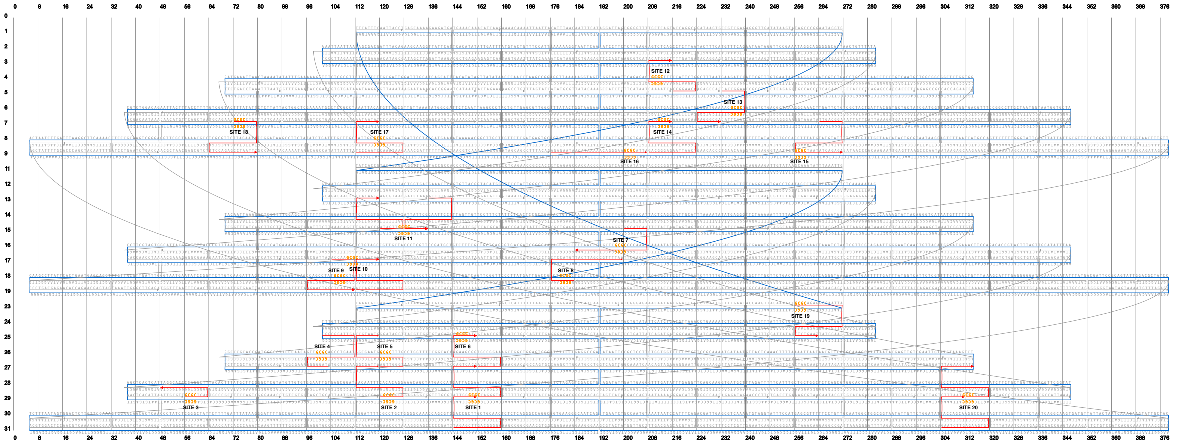

**Supplementary Figure S1:** Cadnano representation of the DNA triangle, with GCGC sites highlighted.

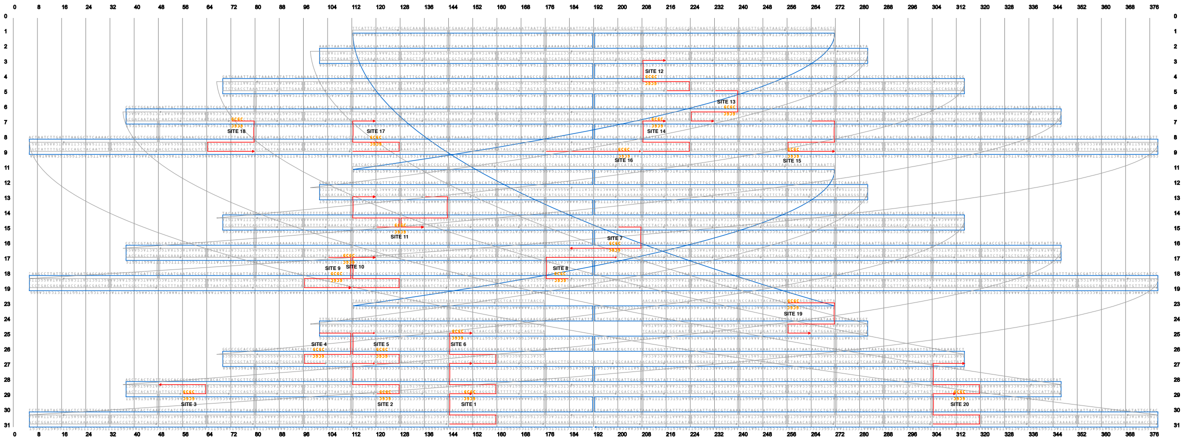

**Supplementary Figure S2:** Same as Fig. S1, but for the four staple deficient (4sd) triangle.

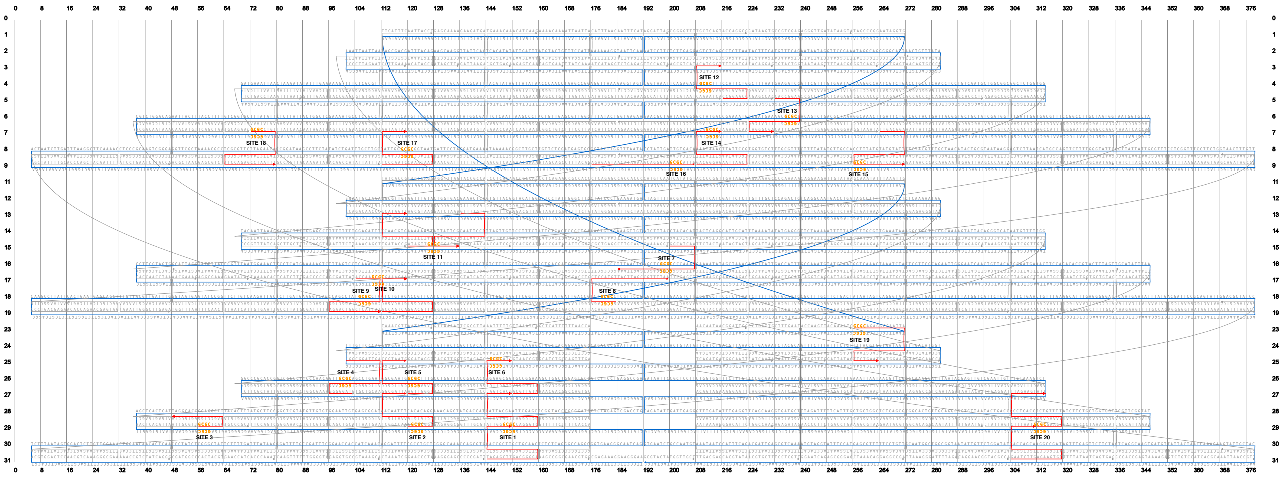

**Supplementary Figure S3:** Same as Fig. S1, but for the eight staple deficient (8sd) triangle.

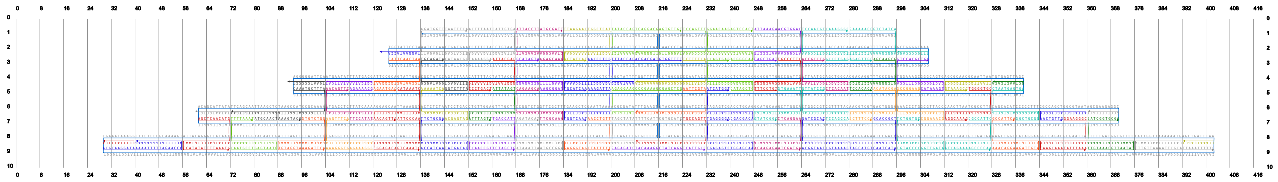

**Supplementary Figure S4:** Cadnano file of DNA isolated trapezoid.

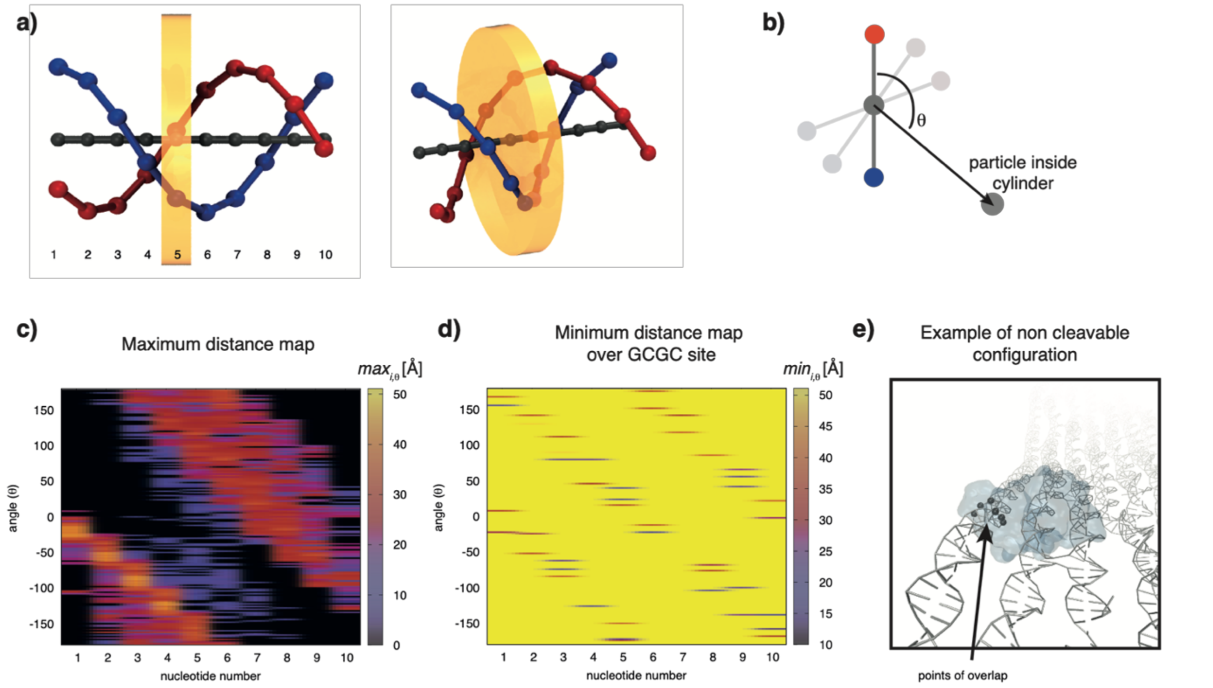


**Supplementary Figure S5:** Geometric criteria used to build the maximum and minimum distance maps. a) A perfect dsDNA of ten base-pairs (bp), used as a reference frame. For each of the ten bp, a cylinder centered on the bp is defined, with the axis parallel to the DNA centerline, and the height equal to the base-to-base distance and infinite radius. Any particle inside the i-th cylinder is considered when constructing the distance maps for the corresponding bp. To compute the angle $\theta$, we use one of the two strands as a reference and compute the angle between its bp vector and the particle position vector (see b). Thus, the angular reference frame rotates along the double strand.  c) Maximum distance map required by the protein ( $max_{i,\theta}$ ) to dock on the DNA. d) Example of minimum distance map ( $min_{i,\theta}$ ) constructed for site 8 (on the scaffold side), for a single frame. In this case, the comparison between the two maps indicates that the site is not accessible; indeed, there is overlap between the protein residues and one of the adjacent strands (see e).

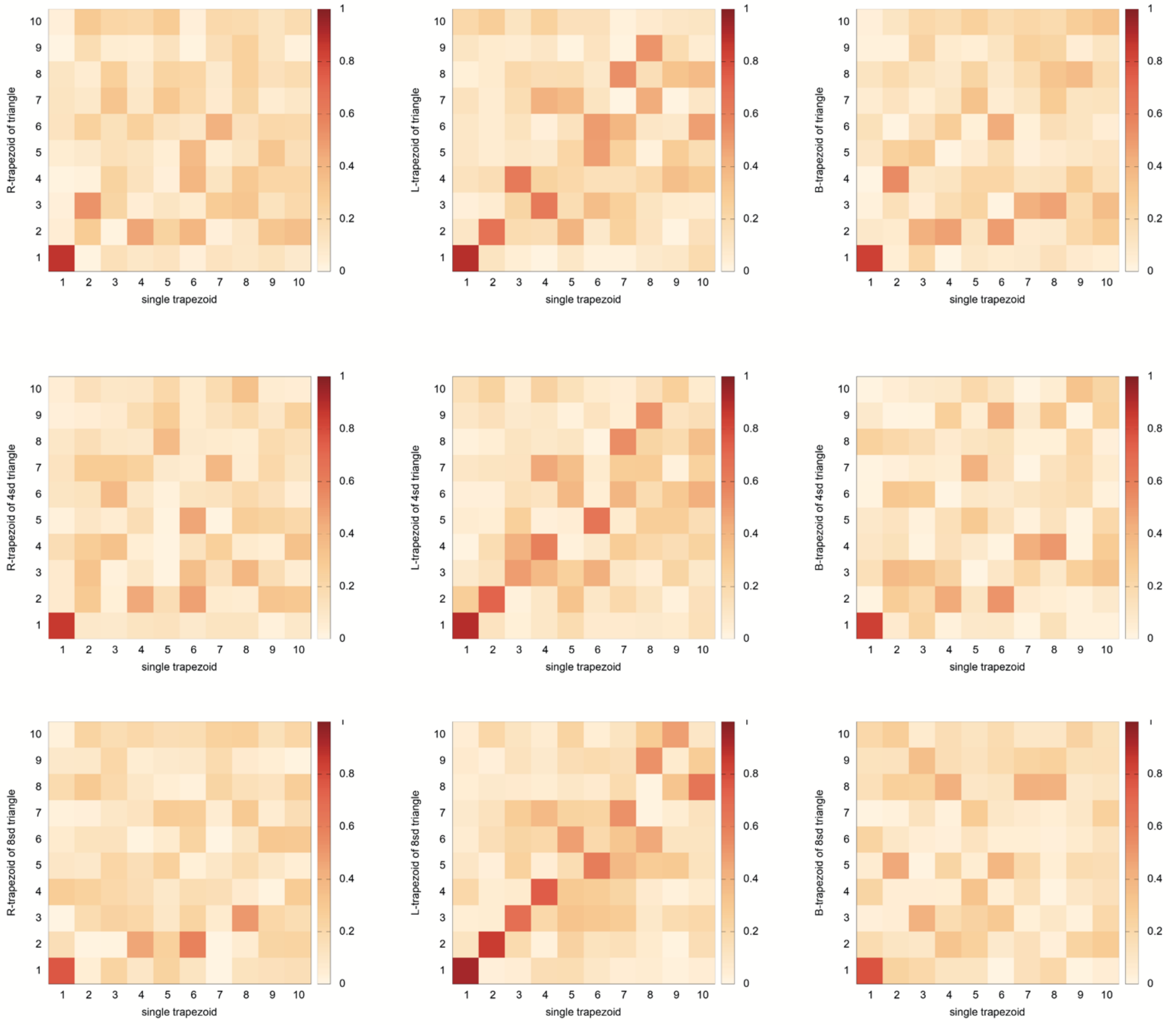


**Supplementary Figure S6:** Absolute value of the inner product between the first 10 eigenvectors of the isolated trapezoid of Fig. 3a and the first 10 eigenvectors of R-, L- and B- trapezoids (first, second and third column) of triangle, 4sd triangle and 8sd triangle (first, second and third row). The first eigenvector direction is conserved across all trapezoids.

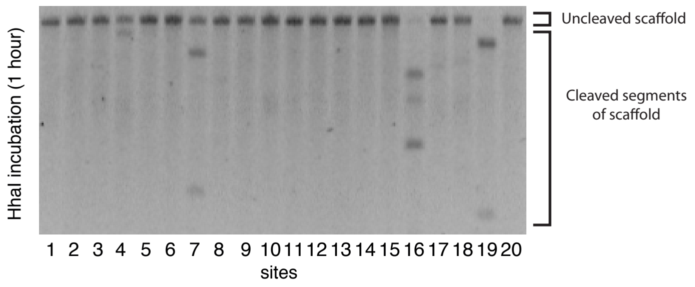


**Supplementary Figure S7:** Gel electrophoretic analysis of the results of ref. [1], showing cleavage of the triangle DNA origami with HhaI after 1 hour incubation. The top gel band represent the fraction of scaffolds not cleaved, while the two faster-electrophoresing bands represent the two segments of the cleaved scaffolds. In order to have an unbiased measurement of the experimentally determined percentage of cut substrate reported in Fig. 3d, we measured, for each band of the gel, the intensity of the highest peak ($i_{un}$), representing uncut scaffolds, and of the two peaks representing the cut scaffold, split in two parts ($i_{c1}$, $i_{c2}$), using the software ImageJ. We then determined the cleavage percentage as: $(i_{c1}+ i_{c2})/(i_{un}+i_{c1}+i_{c2})$. For site 4, where the third band has an intensity below the background noise, the $i_{c2}$ value was assumed, based on the observation that the ratio $i_{c2}/(i_{c1}+i_{c2})$ should be equal to the cleaved scaffold fraction (this is also true for sites 12, 16, 19, within an error of 15%). We assigned to the percent cleavage values an error bar at 20%, using as reference the difference in estimations for the intensities, either with or without the background noise.

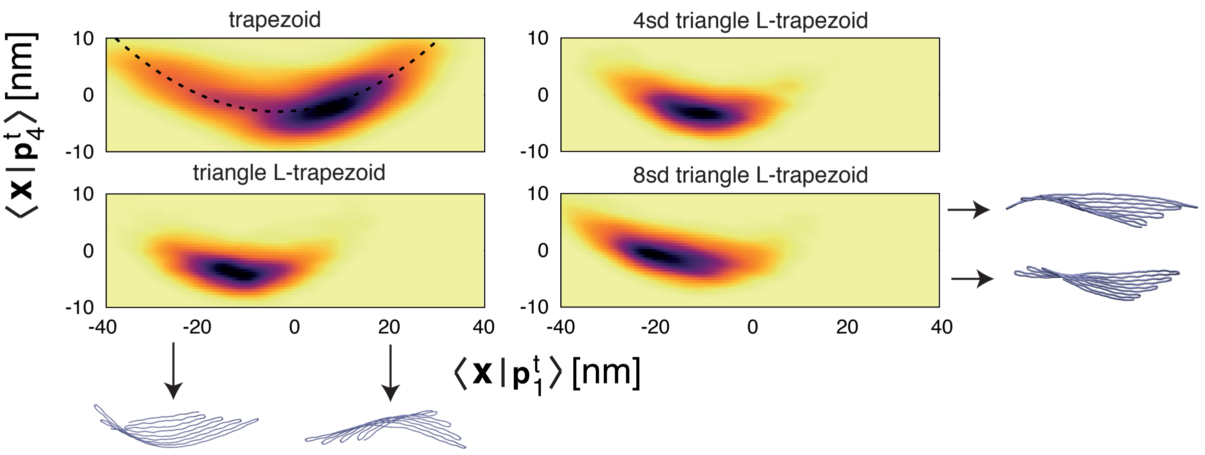


**Supplementary Figure S8:** Two-dimensional histogram of trajectory projected along the first and fourth eigenvector of the isolated trapezoid. The two directions are related through a quadratic law. On the sides, configurations associated to each projection.

**Supplementary Movie S1-S2-S3:** Trajectory of triangle (S1), 4sd triangle (S2), 8sd triangle (S3), corresponding to the time series of Fig. 2a, with configurations colored in blue or red depending on the system being in state 1 or state 2 of Fig. 2b.

**Supplementary Files initialization_files.zip:** Zip file containing lammps configuration files for the three triangles considered and the isolated trapezoid (triangle_input.lammps 4sd_triangle_input.lammps 8sd_triangle_input.lammps trapezoid_input.lammps) and file to execute the 22 September version of LAMMPS (oxdna.lammps).


[1] Stopar, A., Coral, L., Di Giacomo, S., Adedeji, A. F., and Castronovo, M. (2017) Binary control of enzymatic cleavage of DNA origami by structural antideterminants. Nucleic Acids Res., 46 (2), 995–1006.
